# Nuclear tau aggregates inhibit RNA export and form by secondary seeding from cytosolic tau aggregates

**DOI:** 10.64898/2026.02.17.706203

**Authors:** Carolyn J. Decker, Kathleen McCann, Evan Lester, James Pratt, Meaghan Van Alstyne, Yuchen Wang, Roy Parker

## Abstract

Tau aggregates contribute to multiple neurodegenerative diseases including frontotemporal dementia and Alzheimer’s disease (AD). In models of tauopathy and in patient tissue, tau aggregates can form in the cytoplasm, perinuclear region, and nucleus. Using a HEK293T tau biosensor system, we identified that cytoplasmic tau aggregates formed first, followed by perinuclear-ring-like tau assemblies, and then nuclear tau aggregates formed in nuclear speckles. Nuclear tau aggregates only form in cells with pre-existing cytoplasmic tau aggregates and mostly form independently of cells traversing mitosis. Finally, nuclear tau aggregates do not contain exogenous tau seeds and arise by a secondary seeding event dependent on VCP. Nuclear tau aggregates inhibit mRNA export and show a twofold increase in poly-adenylated mRNAs in the nucleus. Together, these findings indicate that nuclear tau aggregation alters RNA biogenesis and occurs by a secondary seeding event from cytoplasmic tau aggregates, which could contribute to tau pathology.

## Introduction

Tau aggregates are insoluble accumulations of tau (MAPT) and associated proteins and RNAs found in the brains of patients with neurodegenerative tauopathies. There are over 20 different tauopathies including Alzheimer’s Disease (AD), frontotemporal dementia (FTD-tau), and Chronic Traumatic Encephalopathy (CTE) [1,2]. Tau fibrillar species can propagate by prion-like mechanisms and can occur in a variety of specific structural forms [3]. Higher order tau assemblies have been shown to be toxic to cells, with smaller soluble aggregates thought to exhibit greater toxicity than large fibrils [4]. How tau aggregates trigger cell toxicity and neurodegeneration in tauopathies is poorly understood.

Cytoplasmic tau aggregates can contain RNA and RNA-binding proteins, including proteins normally localized to nuclear speckles [2,5–7]. Nuclear speckles (also called splicing speckles) are membrane-less organelles that contain nascent RNA polymerase II transcripts, the splicing machinery and RNA export factors [8,9]. SRRM2 (serine/arginine repetitive matrix protein 2) and PNN (pinin), two components of speckles that mislocalize with cytoplasmic tau aggregates [5], contain polyserine domains that are necessary and sufficient for localization to cytoplasmic tau aggregates [2]. At least in cell line models, tau aggregates preferentially grow in association with cytoplasmic assemblies that contain SRRM2 and PNN [2]. These cytoplasmic assemblies can form during mitosis as mitotic interchromatin granules (MIGs) [10] or independently of mitosis where they are referred to as cytoplasmic speckles (CSs) [2]. Polyserine domains can also promote tau aggregation in mice, in human iPSC neurons in culture, and in vitro with recombinant proteins, demonstrating the generality of this effect [11,12].

Several groups have observed that tau can aggregate in the nucleus in cell line models of tau aggregation [5,13,14]. Moreover, nuclear tau assemblies have been observed in postmortem analyses from patients with tauopathies [15–18]. In addition, tau assemblies in association with the nuclear envelope have been described in cell line models and in post-mortem AD samples [16,19]. Where examined, nuclear tau aggregates are exclusively localized to nuclear speckles, perhaps because nuclear speckles are enriched in proteins with polyserine domains [2,5]. The mechanisms that lead to the formation of nuclear tau aggregates in nuclear speckles and their relationship with the formation of cytoplasmic tau aggregates are unknown.

Herein, we use a cell line model of tau aggregation expressing fluorescently tagged versions of tau to address these issues. Upon tau seeding, we discover cytoplasmic tau aggregates form first, followed by formation of nuclear tau aggregates. This suggests that cytoplasmic aggregates are the source for the seeds that induce nuclear tau aggregation. Moreover, nuclear tau aggregates do not contain exogenous seeds and are dependent on VCP function, which can generate new tau seeds [20,21]. Nuclear tau aggregation is dependent on SRRM2 but not PNN, and requires the polyserine domain of SRRM2, which either promotes seed formation, nuclear import of tau seeds, or creates a nuclear speckle environment conducive for tau aggregate propagation. We also observed that specific mRNAs and bulk poly(A)+ RNA accumulate in nuclei with nuclear tau aggregates, demonstrating that tau aggregation within nuclear speckles can inhibit mRNA export. These observations highlight unique cellular perturbations caused by nuclear tau aggregates, which could contribute to toxicity in any tauopathy with nuclear tau aggregates.

## Methods

### Cell culture and tau seeding of HEK293T tau biosensor cells

HEK293T cells stably expressing the 4R RD domain of tau with P301S mutation (ATCC Cat# CRL3275; RRID:CVCL_DA04) were cultured in DMEM with 10% FBS and 1% penicillin-streptomycin antibiotics at 37^°^C with 5% CO_2_ in the Cell Culture Facility RRID: SCR_01898 (CCF). Cells were seeded on poly-L-lysine (Santa Cruz Biotechnology sc-286689A) coated glass coverslips in 24 well plates (Corning 3526) at 125000 cells/well. 24 hours later, a 50µl mix containing 0.875µl 1mg/ml clarified brain homogenate from Tg2541 tau mice (as described in [5]), 9.125µl 1XPBS, 39.25µl DMEM without FBS or antibiotics and 0.75µl lipofectamine 3000 (Thermo Fisher Scientific L300008) per well was prepared, incubated at room temperature for 20 min, then 50µl added to each well. Cells were treated for 24 hours at 37°C then fixed in 4% paraformaldehyde in PBS, washed with PBS, permeabilized with 0.1% Triton X-100, washed with PBS then mounted using Prolong Glass with NucBlue (Thermo Fisher Scientific P36981). Samples were imaged using a Nikon CSU-W1 SoRa confocal spinning disk microscope with a PLAN APO Lambda 100X oil immersion objective lens NA 1.45 and Hammamatsu ORCA Fusion BT sCMOS camera unless otherwise indicated.

### Live cell imaging of HEK293T tau biosensor cells after tau seeding

HEK293T tau biosensor cells were seeded on a poly-L-lysine coated 24 well glass bottom plate (Cellvis P241.5HN) and transfected with clarified tau mouse brain homogenate as described above. Hoeschst 33342 was added to cell culture media to stain nuclei. 6 hours post transfection, the plate was imaged every 15 min for 19.5 hours using the CSU-W1 SoRa confocal spinning disk microscope with an APO LWD 40X NA 1.15 water immersion objective lens, Andor iXon Life 897 EMCCD camera and Okolab Incubation enclosure.

### Purification of 2N4R-Cys-P301S tau

As previously described [11,12], a pET29 plasmid encoding full length 2N4R-Cys-P301S tau was transformed into Rosetta2(DE3)pLysS *e*. *coli*. Expression was induced with 500 µM IPTG for 4 hours with shaking at 37°C. Pellets were collected and lysed in lysis buffer (50 mM Tris pH 7.4, 500 mM NaCl, 1 mM DTT, and 30 mM imidazole, supplemented with one cOmplete ULTRA EDTA-free protease inhibitor tablet (Sigma 6538282001)) via sonication. Lysate was clarified and filtered before added to pre-equilibrated Ni-NTA resin. The column was washed with 5 column volumes (CV) lysis buffer, 10 CV wash buffer 1 (lysis buffer with 200 mM NaCl), then 10 CV wash buffer 2 (lysis buffer with 1M NaCl). Elution buffer (lysis buffer with 300 mM imidazole, 200 mM NaCl) was added, and the column incubated for 30 minutes. Elution was dialyzed overnight into Buffer A (50 mM MES pH 6.00, 50 mM NaCl, 1 mM DTT). Dialyzed protein was then loaded onto a 5 ml Hi-Trap Capto SP ImpRes cation exchange column (Cytiva 17546855) and protein was eluted with a linear gradient of Buffer B (50 mM MES pH6.00, 1M NaCl, 1 mM DTT). Protein containing fractions were pooled and concentrated to 1 mL total volume before loading onto a Superdex 200 pg 16/600 (Cytiva 28989335) column. The column was eluted with 1X PBS pH 7.4, 1 mM TCEP. Protein containing fractions were pooled and concentrated to 50 µM for downstream labeling.

### 2N4R-Cys-P301S maleimide labeling

Purified protein was incubated with 3-fold molar excess of JF549-Maleimide (Tocris 6500) or JF646-Maleimide (Tocris 6590) for 2 hours at room temperature. Excess dye was removed and inactivated with overnight dialysis into 1x PBS pH 7.4, 1 mM DTT, and protein stored at 4°C.

### Exogenous fluorescent seed preparation

5 µM of maleimide-labeled tau was incubated with 40 µg heparin, 1 mM DTT, brought up to 50 µL total volume with fibrilization buffer (1x PBS pH 7.4, 1 mM DTT), and shaken at 37°C for 24 hours. Reactions were pooled and tau fibers were pelleted at 100K xg for 30 minutes. Supernatant was removed and the pellet was resuspended in 1x PBS pH 7.4. Resuspended pellets were sonicated in 15 second pulses with a 15 second recovery for 30 total minutes at an amplitude of 50 using a cup horn attachment on a Fisher Sonic Dismembrator to generate seeds immediately before transfection into HEK293T tau biosensor cells.

### Confocal Microscopy of Exogenous Seeds in HEK293T Tau Biosensor Cells

HEK293T tau biosensor cells were plated on poly-L-lysine (Santa Cruz Biotechnology sc-286689A) coated glass coverslips in 24 well plates (Corning 3526) at 125,000 cells/well. 24 hours later, a 50µl mix containing 3µl sonicated, maleimide-labeled tau seeds, 7µl 1XPBS, 40µl DMEM without FBS or antibiotics and 1µl lipofectamine 3000 (Thermo Fisher Scientific L300008) per well was prepared, incubated at room temperature for 10 min, then 50µl added to each well. For live cell imaging, Hoeschst 33342 was added to cell culture media to stain nuclei. 0.5 hours post transfection, the plate was imaged every 15 min for 19.5 hours using the CSU-W1 SoRa confocal spinning disk microscope with an APO LWD 40X NA 1.15 water immersion objective lens, Andor iXon Life 897 EMCCD camera and Okolab Incubation enclosure. For experiments with a second tau seeding step, cells were transfected after 20 hours with Cy5-maleimide labeled tau seeds as described above. Cells were fixed with 4% PFA after either four hours or twenty-four hours after seed transfection. Cells were then washed once with 1x PBS pH 7.4, permeabilized with 0.5% Triton X-100 for 10 minutes, washed once with 1xPBS with DAPI, and mounted with Prolong Glass. Coverslips were imaged with a Nikon CSU-W1 SoRa confocal spinning disk microscope with a PLAN APO Lambda 100X oil immersion objective lens NA 1.45 and Hammamatsu ORCA Fusion BT sCMOS camera with 0.5 µm z-stacks. Max projections were made using imageJ and used for downstream quantitative analysis. Image analysis was performed using CellProfiler. In short, single cells were segmented by identifying nuclei, and secondary objects of single cells were defined using the tau fluorescence channel and contained one nucleus each. Cy3 tau seeds and tau aggregates were identified in cells. Tau aggregates were then filtered for those that contained a Cy3 tau seed object. Cells were then filtered for those with cytoplasmic or nuclear tau aggregates and further filtered for those with tau aggregates that contained a Cy3 seed.

### RAN Immunofluorescence

HEK293T tau biosensor cells were seeded, transfected with tau mouse brain homogenate, fixed and permeabilized as described above then blocked with 1% BSA 1X PBS for 1 hour, incubated with anti-Ran mouse monoclonal antibody (BD Biosciences 610340) 1:250 in 1% BSA in 1X PBS overnight at 4°C, washed 3 times with 0.1% Tween 20 in 1XPBS, incubated with goat Anti-Mouse IgG AlexaFluor647 (abcam ab150115) 1:1000 in 1% BSA in 1XPBS for 1-2 hours, washed 3 times with 0.1% Tween20 in 1x PBS, then mounted with Prolong Glass with NucBlue. Cell Profiler was used to segment nuclei and cytoplasm of cells, identify cells with nuclear or cytoplasmic tau aggregates and to determine the mean intensity of RAN fluorescence intensity in the cytoplasm and nucleus.

### NLS-mCherry Expression and Nuclear:Cytoplasmic Quantification

HEK293T tau biosensor cells were plated on poly-L-lysine (Santa Cruz Biotechnology sc-286689A) coated glass coverslips at 125000 cells/well. 24 hours later, cells were transfected with plasmid encoding nls-mCherry (nls-mCherry was a gift from Rob Parton (Addgene plasmid # 108881; http://n2t.net/addgene:108881; RRID:Addgene_108881)[22]) using Lipofectamine 3000 (Thermo Fisher Scientific L300008) as per manufacturer’s specification. 24 hours following plasmid transfection, cells were transfected with 1 µL of tau brain homogenate from Tg2541 mice as described above. 24 hours following tau brain homogenate transfection, cells were fixed with 4% PFA and imaged using a Nikon CSU-W1 SoRa confocal spinning disk microscope with a PLAN APO Lambda 100X oil immersion objective lens NA 1.45 and Hammamatsu ORCA Fusion BT sCMOS camera. The nuclear:cytoplasmic ratio was determined for single cells by measuring the total nls-mCherry fluorescence in the nucleus and dividing that value by the total nls-mCherry fluorescence in the cytoplasm. Image analysis was performed using CellProfiler. Single cells were identified by first identifying nuclei, then using the tau fluorescence channel, single cells were identified to contain one nucleus. The cytoplasm and nucleus of single cells were then masked, and the total fluorescence of nls-mCherry was measured behind each mask.

### VCP Inhibitor Treatment and imaging of HEK293T tau biosensor cells

HEK293T tau biosensor cells were seeded on poly-L-lysine (Santa Cruz Biotechnology sc-286689A) coated glass coverslips in 24 well plates (Corning 3526) at 250000 cells/well. 24 hours later, a 50µl mix containing 2µl 1mg/ml clarified brain homogenate from Tg2541 tau mice (as described in [5]), 8µl 1XPBS, 39µl DMEM without FBS or antibiotics and 1µl lipofectamine 3000 (Thermo Fisher Scientific L300008) per well was prepared, incubated at room temperature for 10 min, then 50µl added to each well. 4 hours after tau seeding, the cells were treated with 25μL of VCP inhibitors (2.5μM CB-5083 (MedChem Express HY-12861), 2μM NMS-873 (MedChem Express HY-15713)) or 25μL of 0.5% DMSO in PBS. 20 hours later, the cells were fixed in 4% paraformaldehyde in PBS, washed with PBS, permeabilized with 0.1% Triton X-100, washed with PBS, then mounted using Prolong Glass with NucBlue (Thermo Fisher Scientific P36981). Samples were imaged using a Nikon CSU-W1 SoRa confocal spinning disk microscope with a PLAN APO Lambda 100X oil immersion objective lens NA 1.45 and Hammamatsu ORCA Fusion BT sCMOS camera. The number of non-mitotic cells having cytosolic tau aggregates and nuclear tau aggregates was quantified manually using cell counter in ImageJ. Statisics were performed using the Kruskal-Wallis test followed by Dunn’s Test for Multiple Comparisons.

### SRRM2 enrichment in cytoplasmic tau aggregates

HEK293T tau biosensor cells with at least one allele of SRRM2 fused to Halo tag (previously described in [2]) was simultaneously treated with tau mouse brain homogenate as described above and 200nM Janiela Fluor HaloTag 646 (Promega GA1120) for 24 hours. Cells with nuclear and cytoplasmic tau aggregates were identified, and 1.5 micron square ROIs were drawn in the cytoplasmic aggregate in each cell and in the cytoplasm outside the aggregate (the status of SRRM2 halo signal was unknown when both ROIs were drawn). The integrated density of halo signal in each ROI was measured and the ratio of halo signal in the aggregate over the signal in the cytoplasm was determined as a measure of enrichment.

### SRRM2 and PNN siRNA treatment

HEK293T tau biosensor cells were transfected with 25pmole control siRNA (Thermo Fisher 4390843), SRRM2 siRNA (Thermo Fisher s24003) and/or PNN siRNA (Thermo Fisher s10758) and Lipofectamine RNAiMAX (Thermo Fisher 13778075) in 6 well plates as described and validated in [2]. 24 hours post-transfection, the cells were typsinized and seeded onto poly-L-lysine coated glass coverslips in 24 well plates. 24 hours post-plating, the cells were treated with tau mouse brain homogenate for 24 hours then fixed and permeabilized and immunofluorescence using rabbit anti-SRRM2 antibody (Thermo Fisher PA5-66827) 1:400 or rabbit anti-pinin antibody (ProteinTech 18266-1-AP) 1:200 and anti-rabbit IgG AlexaFluor647 (abcam50079) 1:1000 was performed as described above. The immunofluorescence was used to confirm the effectiveness of the siRNA treatments. Cell Profiler was used to segment nuclei and cytoplasm of cells, identify cells with nuclear or cytoplasmic tau aggregates and to determine the percentage of cells with cytoplasmic aggregates that did or did not have nuclear aggregates.

### SRRM2 DsiRNA treatment

A 27mer duplex DsiRNA directed to sequences between the 25 polyserine and 42 polyserine tracts in the C-terminus of SRRM2 was designed using IDT DsiRNA design tool. A mix containing the duplexed RNA oligos 5’ rGrCrU rCrUrA rGrCrC rUrUrC rCrUrG rUrGrC rArArC rCrUG A 3’ and 5’ rUrCrA rGrGrU rUrGrC rArCrA rGrGrA rArGrG rCrUrA rGrArG rCrCrC 3’ was resuspended in nuclease free H_2_O to 100µM. Negative control DsiRNA (IDT 492561622) and the SRRM2 DsiRNA were diluted in Duplex Buffer (100mM potassium acetate, 30mM Hepes pH 7.5, IDT 493425811). HEK293T tau biosensor cells with an allele of full-length SRRM2 (SRRM2_FL amino acids 1-2748) or SRRM2 in which 205 amino acids have been deleted from the C-terminus (SRRM2_2 amino acids 1-2543), that includes the 25 polyserine and 42 polyserine tracts, fused to Halo tag [2] were transfected in 6 well plates with 10nM SRRM2 DsiRNA or negative control DsiRNA using Lipofectamine RNAiMAX for 24 hours, then typsinized and seeded onto poly-L-lysine coated glass coverslips in 24 well plates. 24 hours post-plating, the cells were treated with 200nM Janiela Fluor HaloTag 646 (Promega GA1120) and mock transfected or transfected with tau mouse brain homogenate for 24 hours as described above, then fixed, permeabilized, and mounted with Prolong Glass with NucBlue. Images of mock transfected cells were analyzed to identify nuclei and determine the mean halo intensity within the nuclei using Cell Profiler. Cell Profiler was used on images transfected with tau seeds to segment nuclei and cytoplasm of cells, identify cells with nuclear or cytoplasmic tau aggregates and to determine the percentage of cells with cytoplasmic aggregates that did or did not have nuclear aggregates.

### FISH analysis

Cy3 and Cy5 oligo(dT) 30mer (IDT) and custom FISH probes to PEG3 and NORAD labelled with Quasar 670 (Stellaris) are described in [23]. Custom FISH probes to ATF3 cDNA were designed using the Stellaris RNA FISH Probe Designer (Biosearch Technologies, Petaluma, CA http://www.biosearchtech.com/stellaris-designer). The ATF3 FISH probes were labelled using 5-propargylamino-ddUTP-ATTO-633 (Axxora JBS-NU-1619-633) and a protocol adapted from [24]. HEK293T tau biosensor cells were seeded on poly-L-lysine glass coverslips for 24 hours, treated with tau mouse brain homogenate for 24 hours then fixed and permeabilized. FISH hybridization was performed following Stellaris protocol for adherent cells (https://biosearchassets.biosearchtech.com/assetsv6/bti_stellaris_protocol_adherent_cell.pdf) and coverslips were mounted with Prolong Glass with NucBlue. FISH samples were imaged using a widefield DeltaVision Elite Deconvolution microscope with an Olympus UPlanSApo 100X/1.40-NA oil objective lens and a PCO Edge sCMOS camera with appropriate filters at room temperature using SoftWoRx Imaging software. Images were analyzed using Imaris Image Analysis Software (Bitplane) (University of Colorado-Boulder, BioFrontiers Advanced Light Microscopy Core).

### Statistical Analysis

All statistical analyses were performed using GraphPad Prism.

## RESULTS

### Cytoplasmic tau aggregates precede nuclear tau aggregation

Following seeding, HEK293T biosensor cells form both cytoplasmic and nuclear tau aggregates, with nuclear tau aggregates co-localizing in nuclear splicing speckles [5], where SRRM2 and PNN normally accumulate [25]. As a first step to understand the relationships between cytoplasmic and nuclear tau aggregates, we examined the relationship between cytoplasmic and nuclear tau aggregates in HEK293T tau biosensor cells [13,26]. In these experiments, we seeded the biosensor cells by transfection with brain extracts from mice expressing the tau P301S mutation (Tg2541 mice). These mice develop transmissible tau aggregates in their brains, which can be transfected into the HEK293T tau biosensor cell line to initiate tau aggregation [13,26].

We observed that 24 hours after tau seed transfection, 24% of the cells had cytoplasmic tau aggregates without nuclear aggregates and 11% had cytoplasmic aggregates with nuclear aggregates (Fig. 1A, B). This indicates that nuclear tau aggregation is not required for cytoplasmic tau aggregates. A striking observation was that all cells with nuclear tau aggregates contained at least one cytoplasmic tau aggregate (Fig. 1B). This result suggests that cytoplasmic tau aggregates form first and are a necessary precondition for the seeding of nuclear tau aggregates.

**Fig. 1.**
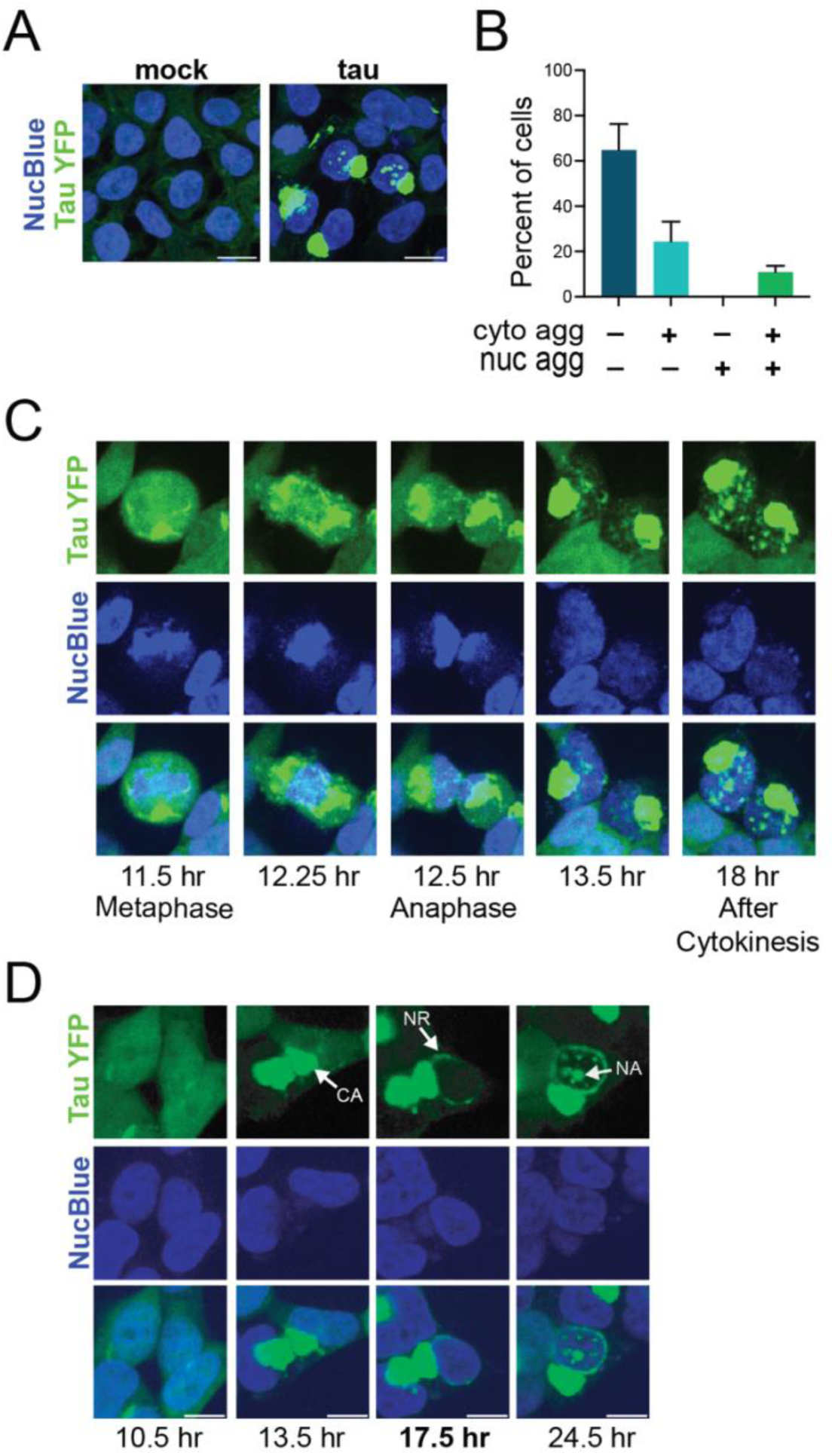
Cytoplasmic tau aggregates precede nuclear tau aggregate formation. (**A**) Tau aggregates observed in the cytoplasm and nuclei of HEK293T tau biosensor cells treated with tau mouse brain homogenate. (**B**) Percentage of HEK293T tau biosensor cells that contain tau aggregates in the cytoplasm or nucleus 24 hours after tau seeding. Mean and SD of 4 independent experiments. (**C-D**) Live cell imaging of HEK293T tau biosensor cells treated with tau mouse brain homogenate. Time post-tau seeding is indicated. (**C**) Images of a cell that undergoes mitosis after cytoplasmic tau aggregates form which is followed by nuclear tau aggregation in the two daughter cells. (**D**) Images of a cell in which tau aggregates form in the nucleus followed by tau assembling around the nucleus and tau aggregation in the nucleus in the absence of mitosis.

To examine directly the temporal relationship between the formation of nuclear and cytoplasmic tau aggregates, we continuously imaged cells starting 6 hours following seeding to monitor the process of tau aggregate formation. We observed two classes of cells that formed nuclear aggregates (Fig. 1C, D). In approximately 20% of the cells, an aggregate formed in the cytoplasm, the cell then underwent mitosis followed by nuclear aggregate formation (Fig. 1C, Movie S1). In these cases, the nuclear tau aggregates may form when tau seeds enter the nucleus during reassembly of the nuclear envelope after mitosis.

In the remaining 80% of cells with nuclear tau aggregates, we observed three phases of tau aggregation (Fig. 1D, Movie S2, S3). First, cytoplasmic tau aggregates (CA) formed and coalesced into large aggregates. Second, perinuclear tau assemblies formed that encircled the exterior of the nucleus forming a nuclear ring (NR). Third, nuclear tau aggregates (NA) appeared. Thus, in the majority of cells, nuclear tau aggregates form after cytoplasmic tau aggregates, independent of mitosis, with an intermediate stage wherein a perinuclear ring of tau assemblies occurs.

To rule out the possibility of a prior unobserved mitosis allowing seeds to enter the nucleus, we imaged cells continuously immediately following tau seeding. We still observed cells that formed nuclear aggregates without undergoing cell division (Movie S4), confirming that a mitotic event is not required for nuclear tau aggregation. The exact nature of the perinuclear ring of tau is unclear but might be related to the coiled body assemblies of tau seen in oligodendrocytes [27–29] or other perinuclear tau assemblies observed in neurons of diseased human brains [16,19,30]. Interestingly, the perinuclear ring of tau shows different dynamic properties from cytoplasmic or nuclear aggregates, and therefore may be a different type of tau assembly [19].

### Nuclear tau aggregation does not correlate with disruptions of nuclear-cytoplasmic transport

The observation that nuclear tau aggregate formation is dependent on prior formation of cytoplasmic aggregates suggests that cytoplasmic aggregates are required for new seeding events in the nucleus. One possible mechanism by which tau seeds could enter the nucleus is that cytoplasmic tau aggregates lead to defects in proper nuclear cytoplasmic transport or nuclear membrane integrity, which then would allow aberrant entry of tau seeds into the nucleus. This possibility is suggested by cytoplasmic protein aggregates interfering with both nuclear import and export [31], mislocalization of the transport factor RAN in transgenic mice with tau aggregates [30], and Nup98 and other phenylalanine-glycine (FG) rich nucleoporins that form a diffusion barrier within nuclear pore complexes co-aggregating with tau and accumulating in NFTs of AD brains [30].

To examine if cytoplasmic tau aggregates alter nuclear cytoplasmic transport in this context, we measured the nuclear to cytoplasmic ratio of RAN by immunofluorescence in HEK293T tau biosensor cells. The nuclear to cytoplasmic gradient of RAN is an indicator of properly controlled transport of proteins and other biomolecules in and out of the nucleus [32–35]. We did not observe a difference in the nuclear to cytoplasmic ratio of RAN in cells with cytoplasmic tau aggregates compared to cells without aggregates (Fig. S1A, B). This result indicates that the aggregation of tau in the cytoplasm does not lead to a general perturbation of transport into the nucleus.

Additionally, we tested if cytoplasmic or nuclear tau aggregates perturb nuclear envelope integrity or nuclear import. To do this, we expressed an NLS-mCherry, which localizes to the nucleus in healthy, non-seeded cells (Fig. S2A, top panel), and calculated the nuclear:cytoplasmic ratio of NLS-mCherry fluorescence in cells with and without tau aggregates. We observed that the nuclear:cytoplasmic ratio of NLS-mCherry did not change in cells with only cytoplasmic aggregates, or in cells with nuclear and cytoplasmic aggregates, compared to cells without aggregates (Fig. S2A, B). Thus, neither cytoplasmic nor nuclear tau aggregates effect the nuclear envelope integrity or nuclear import in the context where nuclear tau aggregates form.

### Secondary seeding events lead to nuclear tau aggregates

Nuclear tau aggregation could occur in at least two non-mitotic ways. Either exogenous seeds translocate into the nucleus and directly seed nuclear aggregates, or cytosolic aggregates are processed to produce a new seeding competent species that subsequently translocate to the nucleus. To test these possibilities, we transfected cells with Cy3-labeled tau seeds to trigger endogenous tau aggregation, fixed cells after 24 hours, and used confocal microscopy to determine the subcellular localization of exogenous seeds relative to cytoplasmic and nuclear tau aggregates (Fig. 2A).

**Fig. 2.**
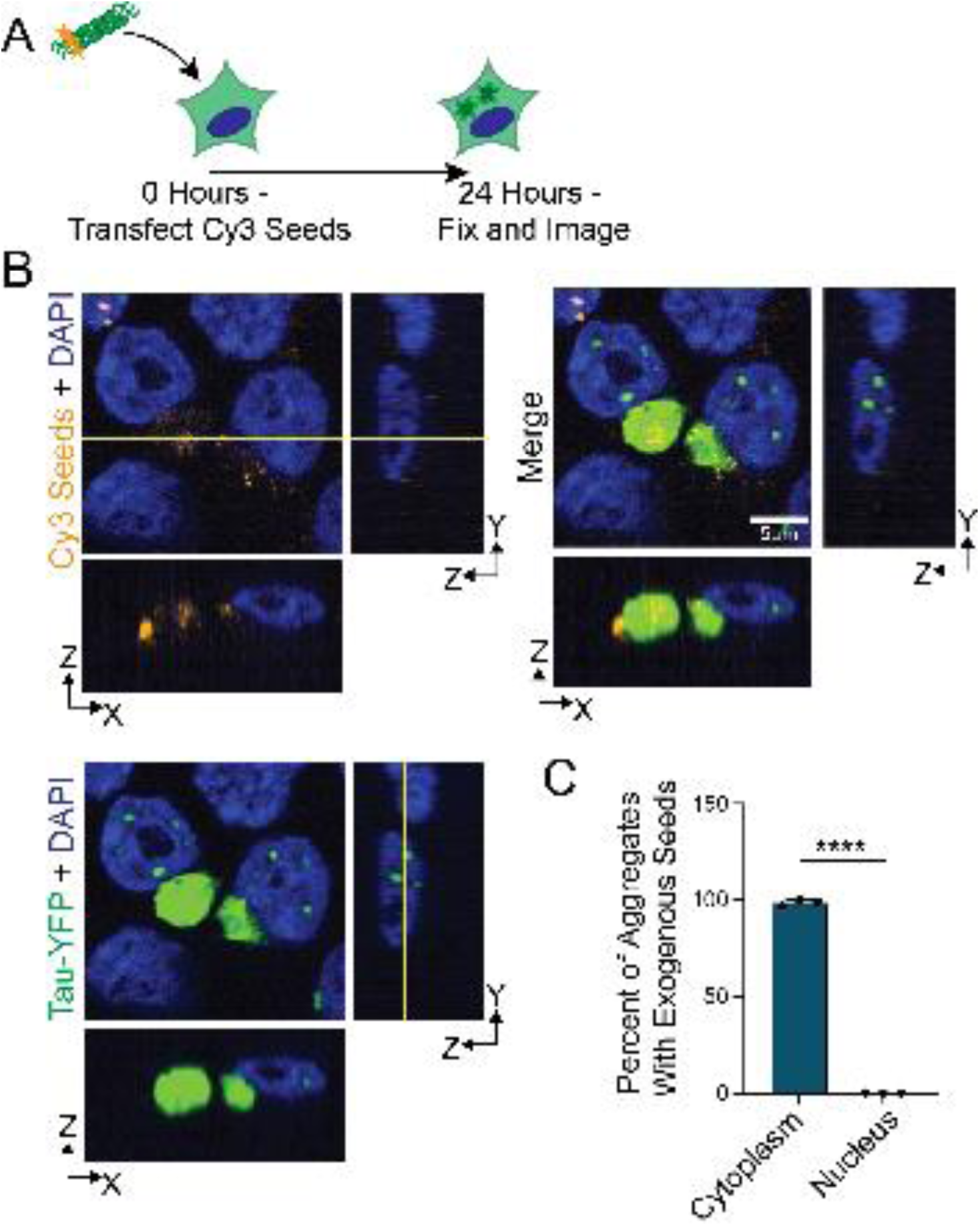
Nuclear tau aggregates do not contain exogenous tau seeds. (**A**) Experimental schematic where fluorescently labeled tau seeds are transfected into HEK293T tau biosensor cells and fixed for confocal microscopy 24 hours later. (**B**) Representative images of fluorescent tau seeds that were transfected into HEK293T tau biosensor cells. Square images are a single Z plane in XY. YZ and XZ projections are shown to the right and below for each channel. (**C**) Quantification of the fraction of cytoplasmic or nuclear tau aggregates that colocalize with exogenous seeds. N = 3 independent experiments, statistics performed with Welch’s T test, p<0.0001.

An important result was that no nuclear tau aggregates colocalized with exogenous Cy3 tau seeds, suggesting that nuclear tau aggregates arise by intracellularly produced seeds. In contrast, essentially all cytoplasmic tau aggregates contain Cy3 fluorescence (Fig. 2B, C). It may be that Cy3 seeds are not able to enter the nucleus as they are sequestered to the cytoplasmic tau aggregates, which always precede nuclear tau aggregation. To address this possibility, we performed a similar experiment in which cells were first transfected with Cy3-labeled tau seeds, then transfected with Cy5-labeled tau seeds 20 hours later, and fixed 24 hours after the initial Cy3 seed transfection (Fig. S3A, B). Here, if exogenous tau seeds can localize to the nucleus after the formation of cytoplasmic aggregates, the Cy5 signal would be localized to nuclear tau aggregates. However, we did not observe Cy5 seeds colocalizing with nuclear tau aggregates or entering the nucleus at all (Fig. S3B). This observation supports the conclusion that new seeds produced intracellularly seed nuclear tau aggregates.

A final formal possibility is that exogenous Cy5 seeds were not able to trigger tau aggregation within 4 hours of transfection. To test this possibility, we transfected Cy5 seeds and fixed cells after 4 hours (Fig. S3C). While we observed small cytoplasmic aggregates that colocalize with Cy5 seeds at this time point, indicating that four hours is sufficient to trigger tau aggregation, we did not observe nuclear tau aggregates (Fig. S3D). Therefore, exogenous seeds do not directly trigger the formation of nuclear tau aggregates suggesting that new seeds produced from cytoplasmic tau aggregates trigger the formation of nuclear tau aggregates.

Tau seeds can be produced from tau aggregates by the action of VCP, which is thought to extract ubiquitinated tau from fibrils and thereby generate smaller seeding competent tau assemblies [20,21]. The role of VCP in generating new tau seeds predicts that VCP function should promote the formation of nuclear tau aggregates. To test this prediction, we inhibited VCP with small molecule inhibitors following tau seeding and then examined the formation of cytoplasmic and nuclear tau aggregates (Fig. 3A).

**Fig. 3.**
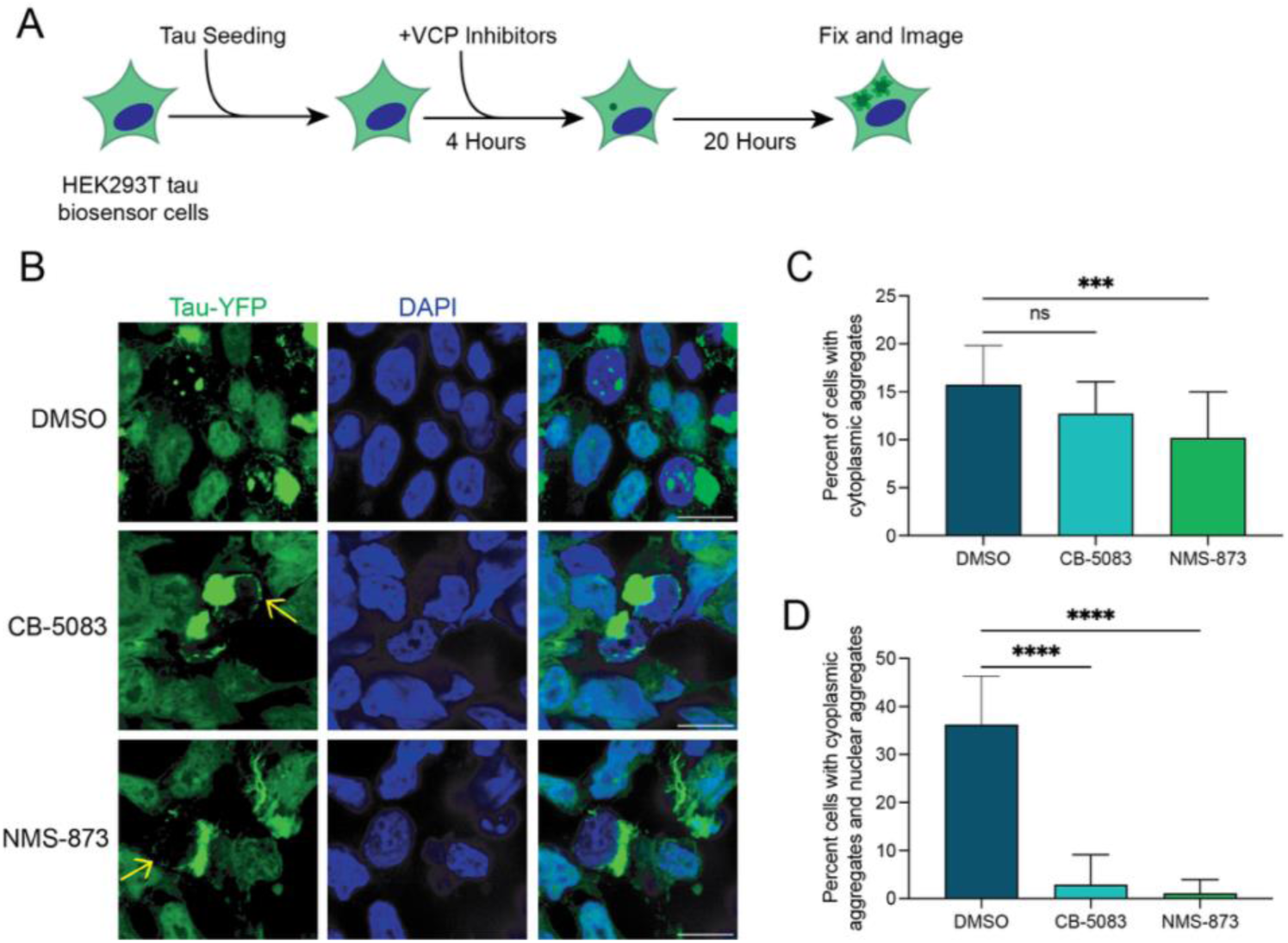
VCP activity is required for the formation of nuclear tau aggregates. (**A**) Experimental design of VCP inhibitor treatment of HEK293T tau biosensor cells. (**B**) Representative confocal microscopy images for each of the treatment groups. Yellow arrows indicate perinuclear tau aggregates. (**C**) Quantification of the percentage of cells having cytosolic tau aggregates. N = 3 independent experiments, statistics performed using the Kruskal-Wallis test followed by Dunn’s Test for Multiple Comparisons. ns = not significant, *** p = 0.0007. (**D**) Quantification of the proportion of cells with cytosolic aggregates that also have nuclear tau aggregates. N = 3 independent experiments, statistics performed using the Kruskal-Wallis test followed by Dunn’s Test for Multiple Comparisons. ns = not significant, **** p < 0.0001.

Strikingly, we observed that inhibition of VCP with CB-5083, which inhibits the ATPase activity of VCP [36], or NMS-873, an allosteric inhibitor [37], both strongly reduced the formation of nuclear tau aggregates in cells that had cytosolic tau aggregates (Fig. 3B-D). We observed no effect on cytosolic tau aggregation with CB-5803. However, while treatment with NMS-873 mildly decreased cytoplasmic tau aggregation as seen earlier (Fig. 3C) [21], the reduction in nuclear tau aggregation was striking and not simply due to the reduction in cytoplasmic aggregates (Fig. 3D). The requirement of VCP function for nuclear tau aggregation argues VCP is responsible for the formation of secondary seeds from cytosolic tau aggregates that can then enter the nucleus and initiate nuclear tau aggregation.

Interestingly, although both CB-5803 and NMS-873 reduced nuclear tau aggregation, we still observed the formation of perinuclear tau assemblies (Fig. 3B, yellow arrows). This demonstrates the formation of the perinuclear ring of tau is independent of VCP activity.

### Nuclear tau aggregates can be dependent on SRRM2

We have previously shown that multiple components of nuclear speckles, including SRRM2, PNN, and MSUT2, can mis-localize to cytosolic tau aggregates [5]. One hypothesis is that these nuclear speckle components interact directly or indirectly with tau in the cytoplasmic aggregates and play a role in localizing tau seeds from the cytosol to the nucleus. To explore this possibility, we first examined whether the presence of nuclear tau aggregates was correlated with the accumulation of SRRM2 in cytoplasmic tau aggregates.

We observed SRRM2 enrichment in cytosolic tau aggregates in all cells with nuclear tau aggregates (Fig. 4A, B), although the degree of enrichment varied between cells (Fig. 4B). This demonstrates that SRRM2 association with cytoplasmic tau aggregates is positively correlated with the formation of nuclear tau aggregates.

**Fig. 4.**
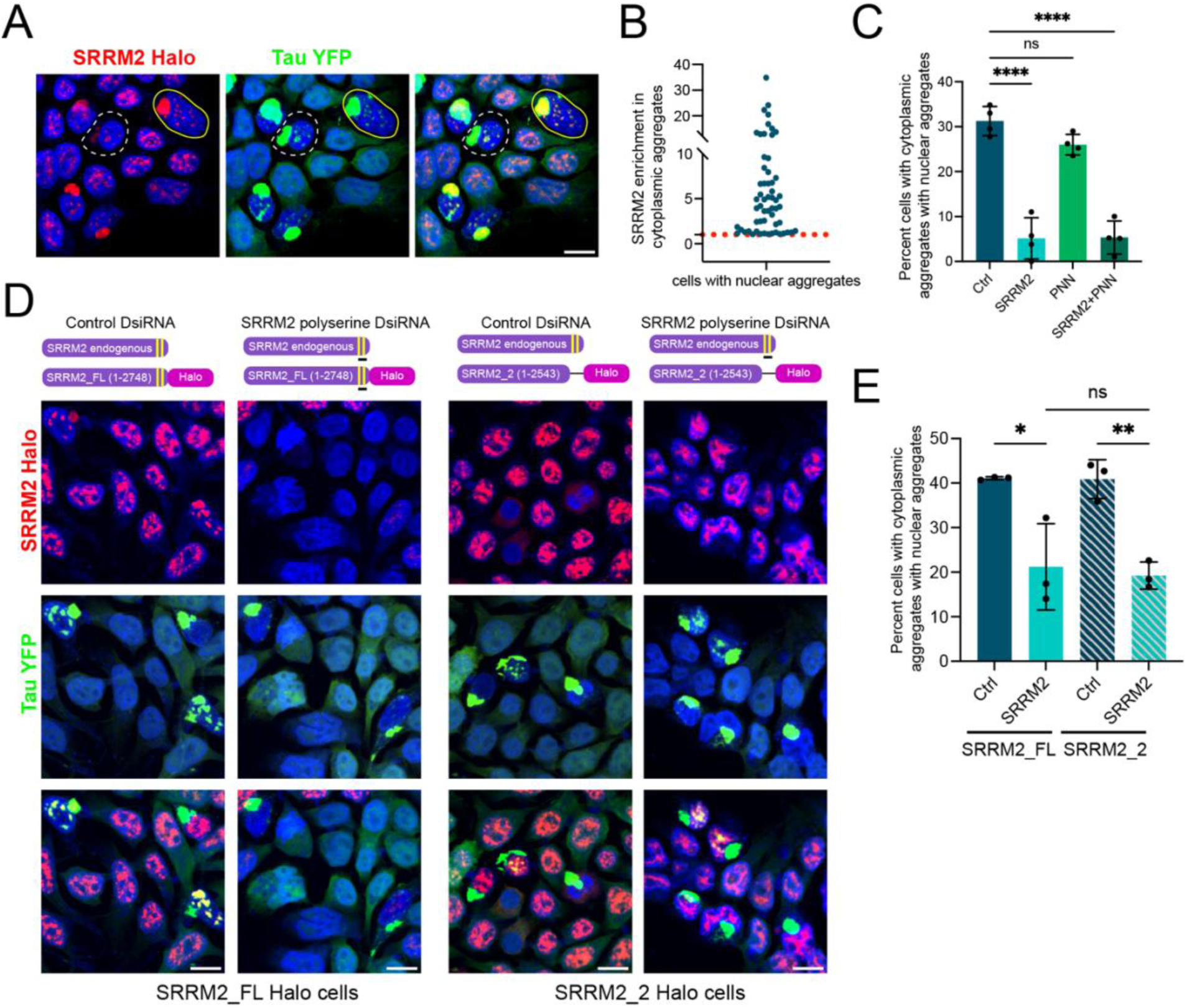
Tau aggregate formation in the nucleus depends on the polyserine domain of nuclear speckle protein SRRM2. (**A**) HEK293T tau biosensor with full-length SRRM2 fused to Halo tag treated with tau mouse brain homogenate for 24 hours. (**B**) Analysis of the enrichment of SRRM2 halo signal in cytoplasmic tau aggregates compared to the cytoplasm in cells that have nuclear tau aggregates. The red dotted line denotes an enrichment of 1. The blue dots represent individual cells. (**C**) Percentage of cells with cytoplasmic aggregates with nuclear aggregates after treatment with SRRM2 or PNN siRNA. N = 4 independent experiments, statistics performed using ordinary one-way ANOVA and Dunnett’s multiple comparisons test, **** p = < 0.0001. (**D**) Illustration of the endogenous SRRM2 allele, full-length SRRM2 (SRRM2_FL) halo tagged allele and halo tagged SRRM2_2 allele which lacks the two C terminal polyserine sequences (yellow bars) and the location of the DsiRNA directed to sequences located between the polyserine sequences (black bar). Images of SRRM2_FL and SRRM2_2 HEK293T tau biosensor cells treated with tau mouse brain homogenate. (**E**) Percentage of SRRM2_FL and SRRM2_2 cells with cytoplasmic aggregates and nuclear aggregates after DsiRNA treatment. Bars for all plots represent mean and SD of 3 independent experiments. N = 3 independent experiments, statistics performed with ordinary one-way Anova and Dunnett’s multiple comparisons test, ns = not significant, * p = 0.0135, ** p = 0.0077.

One possibility is that nuclear speckles, because they are enriched in the polyserine-containing proteins, SRRM2 and PNN, promote tau aggregation similar to cytoplasmic assemblies of these proteins [2]. To test this possibility, we examined how knock-down of SRRM2 or PNN affected nuclear tau aggregation. We have previously found, using a flow cytometry assay to detect tau aggregation in the HEK293T tau biosensor cells via FRET, that knock down of PNN, which is expressed at higher levels than SRRM2, reduced tau aggregation whereas depletion of SRRM2 had no effect [2]. The FRET flow cytometry assay most likely assesses primarily cytoplasmic tau aggregation since the contribution of the smaller, nuclear tau aggregates to the total FRET signal in a cell would be minimal compared to the cytoplasmic aggregates. Given this, we examined how knock-down of SRRM2 or PNN affected nuclear tau aggregation by imaging individual cells and counting the frequency of tau aggregation in the nucleus and cytoplasm. Because nuclear tau aggregation is strongly dependent on the presence of cytoplasmic aggregates, to assess specifically whether knock down of PNN and/or SRRM2 affected nuclear tau aggregation, we only analyzed cells with cytoplasmic aggregates to determine if the frequency of formation of nuclear aggregates was altered.

We find that knockdown of SRRM2 reduced the frequency of nuclear aggregates in cells with cytoplasmic aggregates (Fig. 4C). Knockdown of PNN alone did not significantly reduce nuclear tau aggregation and depletion of both SRRM2 and PNN did not further lower the frequency of nuclear aggregates forming in cells with cytoplasmic aggregates (Fig. 4C). These results indicate that SRRM2 plays a unique role in the aggregation of tau in the nucleus.

To examine if this role was dependent on the polyserine domain of SRRM2, we determined whether cells that only express a form of SRRM2 that lacks the polyserine sequences can form nuclear tau aggregates. We previously used CRISPaint to engineer a polyclonal cell line where a Halo tag was added to at least one allele of endogenous SRRM2 gene [2]. Deleting a region of the C-terminus of SRRM2, which includes both the 25 and 42 amino acid tracts of polyserine, disrupts SRRM2’s localization to cytoplasmic tau aggregates without affecting its localization to nuclear speckles [2]. We specifically depleted the wild-type endogenous SRRM2 protein in the HEK293T tau biosensor cells expressing the SRRM2 Halo protein lacking the polyserine domain by treating the cells with an DsiRNA directed to the deleted region of SRRM2. We used a 27mer DsiRNA to knockdown the endogenous SRRM2 mRNA because standard siRNA prediction software did not find good candidate sequences in the deleted region of SRRM2.

To monitor the effectiveness of the knockdown of full-length SRRM2, we examined the ability of the DsiRNA to reduce the expression of full-length Halo-tagged SRRM2 (Fig. 4D, Fig. S4). SRRM2 DsiRNA treatment of the HEK293T tau biosensor cells with the two full-length SRRM2 alleles strongly reduced the number of Halo positive cells (Fig. 4D, Fig. S4), and, as seen with the previous SRRM2 siRNA treatment, reduced the formation of nuclear tau aggregates (Fig. 4E).

As expected, SRRM2-Halo positive cells are present after SRRM2 DsiRNA treatment of the cells expressing the Halo-tagged SRRM2 polyserine deletion protein (Fig. 4D), although depletion of the full-length SRRM2 protein did slightly reduce the levels of the truncated SRRM2 protein (Fig S4). The DsiRNA treated mutant cells expressing only SRRM2 protein lacking the polyserine domain had a similar defect in nuclear tau aggregation as the cells that were depleted of the full-length SRRM2 protein (Fig. 4E). This result indicates that the polyserine domain of SRRM2 is required for SRRM2 to promote tau aggregation in the nucleus.

The polyserine rich domain of SRRM2 may create a biochemical environment in nuclear speckles that is conducive for tau aggregation as has been observed for cytoplasmic aggregates. Alternatively, the association of SRRM2 with tau aggregates in the cytoplasm, which is dependent on its polyserine domain, could be important for the subsequent seeding or formation of tau aggregates in the nucleus.

### Poly(A)+ RNA accumulates in nuclei with nuclear tau aggregates

Nuclear speckles are thought to play a role in mRNA nuclear export since nuclear export factors of mRNAs are enriched in nuclear speckles [38], and mRNAs that lack introns traffic through nuclear speckles for export [39]. Given this, we examined whether RNA export is altered in cells with nuclear tau aggregates in nuclear speckles by quantifying the amount of nuclear poly(A)+ RNA in cells with or without nuclear tau aggregates.

We observed a 2-fold increase in the total intensity of poly(A)+ RNA in nuclear foci in cells with nuclear tau aggregates (Fig. 5A, E) compared to cells without any tau aggregates or compared to cells with cytoplasmic tau aggregates but no nuclear aggregates. The poly(A)+ RNA accumulation in the nuclear tau aggregates is consistent with changes in the properties or function of nuclear speckles leading to the retention of mRNAs.

**Fig. 5.**
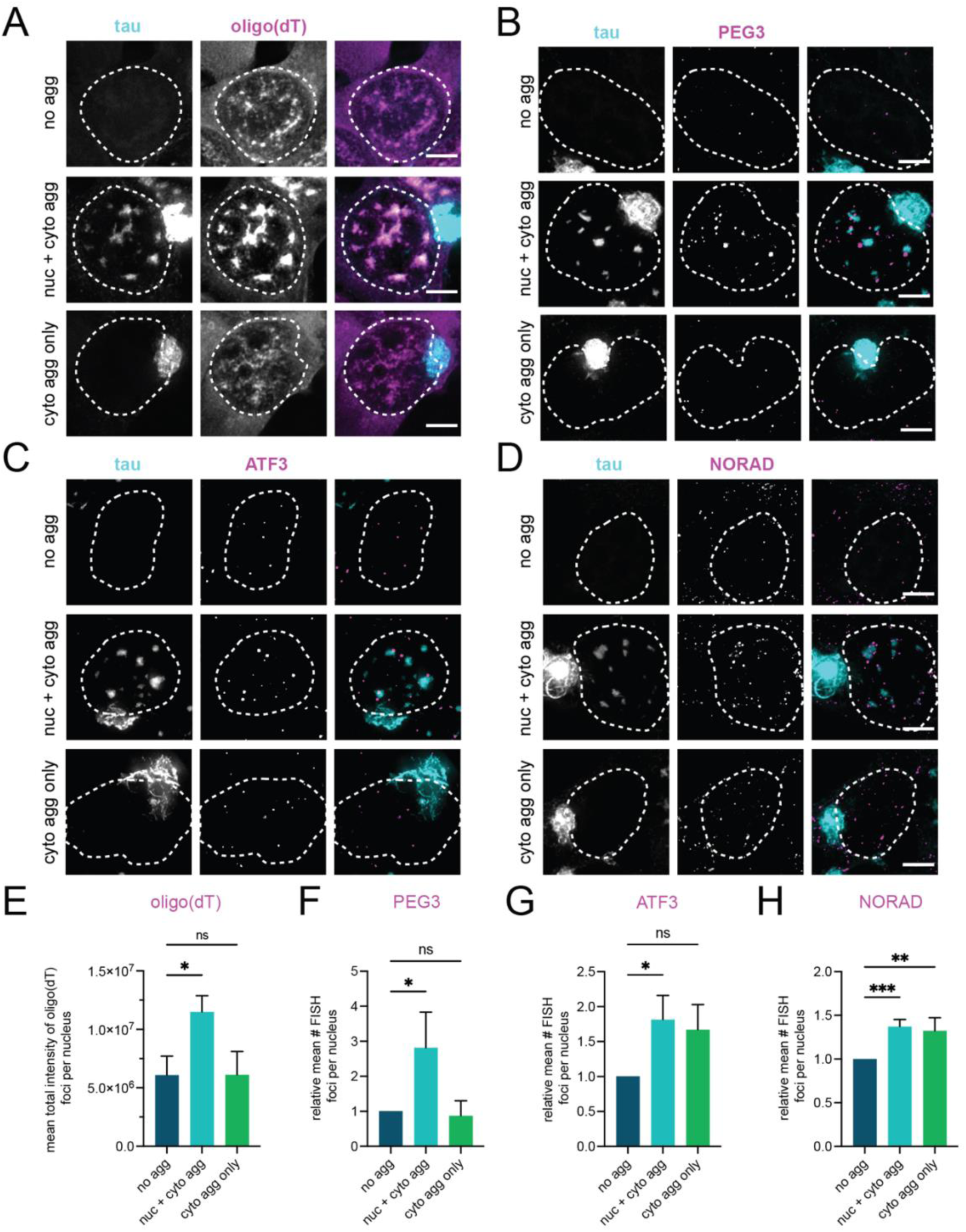
Tau aggregation in the nucleus is associated with the accumulation of polyA+ RNAs in the nucleus. (**A-D**) Images of HEK293T tau biosensor cells treated with tau mouse brain homogenate without any tau aggregates (no agg), with only cytoplasmic tau aggregates (cyto agg only) or with both cytoplasmic and nuclear tau aggregates (nuc + cyto agg). Outline of nucleus indicated by dashed white lines. (**A**) FISH with oligo(dT). (**B**) FISH with probes against PEG3 mRNA. (**C**) FISH with probes against ATF3 mRNA. (**D**) FISH with probes against NORAD ncRNA. (**E)** The mean total intensity of oligo(dT) FISH signal per nucleus in cells with only cytoplasmic aggregates or with both nuclear and cytoplasmic aggregates relative to the mean total intensity observed in cells with no aggregates. Bars for all plots represent mean and SD. N = 3 independent experiments, statistics performed with ordinary one-way Anova and Dunnett’s multiple comparisons test, ns = not significant, * p = 0.0136**. (F-H**) The mean number of FISH foci for the indicated RNAs in cells with only cytoplasmic aggregates or with both nuclear and cytoplasmic aggregates relative to the mean number observed in cells with no aggregates. Bars represent mean and SD. (**F**) The mean number of PEG3 FISH foci. N = 3 independent experiments, statistics performed with ordinary one-way Anova and Dunnett’s multiple comparisons test, ns = not significant, * p = 0.0231. (**G**) The mean number of ATF3 FISH foci. N = 3 independent experiments, statistics performed with ordinary one-way Anova and Dunnett’s multiple comparisons test, ns = not significant, * p = 0.0241. (**H**) The mean number of NORAD FISH foci. N = 4 independent experiments, statistics performed with ordinary one-way Anova and Dunnett’s multiple comparisons test, ** p = 0.0026, *** p = 0.0010.

We also observed that the polyA+ FISH signal in speckles does not directly overlap with tau and instead forms a periphery around the tau aggregate (Fig. S5). This is consistent with earlier data that nuclear tau aggregates alter the spatial organization of nuclear speckles [5]

We also examined specific RNAs, including mRNAs that undergo splicing (PEG3 and ATF3 mRNA) and an unspliced non-coding RNA (NORAD), using smFISH to determine if they accumulated in nuclei containing nuclear tau aggregates. The number of PEG3 FISH spots in nuclei was approximately 2-fold higher in cells with nuclear tau aggregates compared to cells with no tau aggregates or tau aggregates solely in the cytoplasm (Fig. 5B, F). The PEG3 smFISH foci tended to be larger in the cells with nuclear aggregates and localized within close proximity to the tau aggregates (Fig. 5B), similar to the distribution of poly(A)+ RNA, suggesting that nascent PEG3 transcripts are clustering with other RNAs in association with nuclear tau aggregates.

The number of nuclear ATF3 and NORAD FISH spots increased in cells with nuclear aggregates and in cells that only had cytoplasmic aggregates compared to the cells that did not have any aggregates (Fig. 5C, D, G, H). This observation suggests that tau aggregation in the cytoplasm may lead to increased expression of some RNAs. Similar to what we observed with PEG3 mRNA, NORAD FISH spots tended to be larger in cells with nuclear aggregates and localize in proximity to the aggregates (Fig. 5B, D, Fig. S6). There is a significant increase in the volume of NORAD FISH spots, specifically in cells with nuclear aggregates (Fig. S6), suggesting that in addition to increased expression in response to tau aggregation in the cytoplasm, there is retention of NORAD RNA in cells with tau aggregates in nuclear speckles. The increase in nuclear PEG3 mRNA and NORAD supports the interpretation that tau aggregation within nuclear speckles interferes with nuclear export of RNA. RNAs may be physically trapped within aggregates that form in nuclear speckles similar to the lack of exchange of nuclear speckle proteins in nuclear tau aggregates [5].

## Discussion

We present several lines of evidence that nuclear tau aggregates form from, and are dependent on, cytoplasmic tau aggregates. First, every cell with nuclear tau aggregates contains a cytoplasmic tau aggregate (Fig. 1). Second, movies of the process of tau aggregation demonstrate that nuclear tau aggregates form after cytoplasmic tau aggregates (Fig. 1, Movies S1-4). Third, while fluorescent tau seeds can be observed in cytoplasmic tau aggregates, exogenous seeds are not observed in nuclear tau aggregates (Fig. 2). Fourth, inhibition of VCP, which is known to fragment tau fibers to produce seeding competent tau [20,21], reduces the formation of nuclear tau aggregates (Fig. 3). Taken together, this suggests cytoplasmic tau aggregates can generate secondary tau seeds, which can enter the nucleus and initiate the formation of nuclear tau aggregates (Fig. 6).

**Fig. 6.**
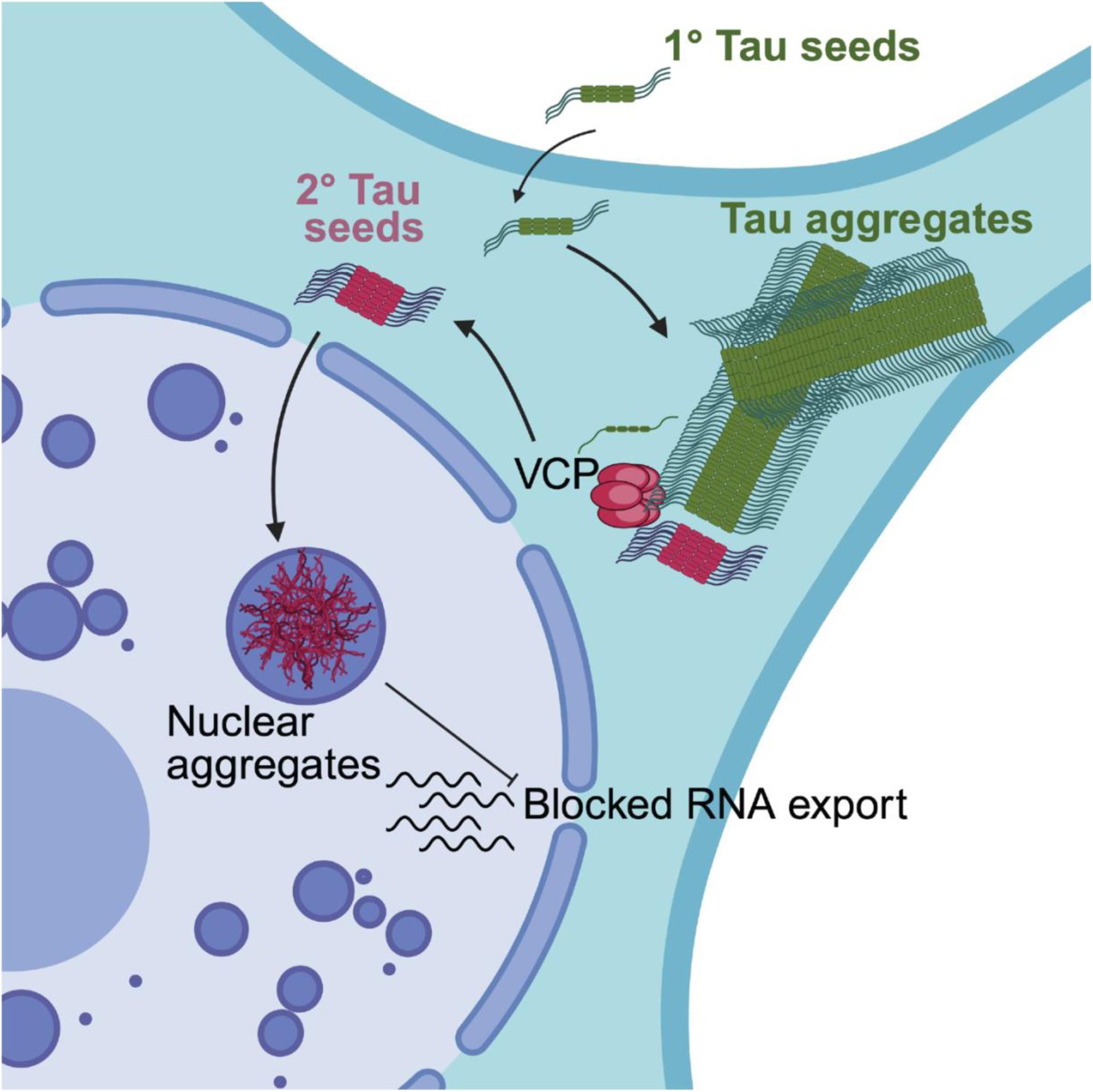
Nuclear tau aggregates form by secondary seeding from cytosolic tau aggregates and inhibit RNA export. Cytosolic tau aggregates form first after a primary (1°) seeding event. Cytolosic tau aggregates are acted upon by VCP, which extracts tau fibrils, thereby breaking the tau fiber and generating seeding competent assemblies (secondary (2^°^) seeds). These 2^°^ seeds enter the nucleus and form nuclear tau aggregates within nuclear speckles. The presence of nuclear tau aggregates disrupts nuclear export and leads to the accumulation of RNAs in the nucleus.

Our data suggest that VCP can generate tau seeds capable of additional seeding within a cell. The key observation is that the formation of nuclear tau aggregates is dependent on VCP (Fig. 3), which is consistent with prior work showing VCP can extract tau from fibrils to generate seeding competent assemblies [20,21]. VCP utilizes several adaptors, which can have different effects on bulk tau aggregation [21]. However, it remains unclear what VCP adaptor(s) are utilized for the formation of nuclear seeding competent tau assemblies.

An unresolved issue is how secondary tau seeds enter the nucleus. In dividing cells, tau seeds could enter the nucleus following mitosis during nuclear reformation, which we observe in ∼20% of the cells with nuclear tau aggregates (Fig. 2). However, in 80% of the cells with nuclear tau aggregates, tau aggregates appear in the nucleus without the cells undergoing a mitosis after the formation of a cytoplasmic tau aggregate (Fig. 2). One possibility is that since cytoplasmic tau aggregates can lead to nuclear envelope dysfunction, damage to the nuclear envelope can allow nuclear tau seed entry by passive diffusion. However, this is unlikely as we do not observe a disruption in the RAN gradient (Fig. S1) or nuclear envelope integrity (Fig. S2). Alternatively, secondary seeds produced from cytoplasmic aggregates could interact with nuclear import machinery, either directly, because of a specific post-translational modification [40], or by binding nuclear localized proteins such as SRRM2, and then be actively imported into the nucleus. This latter model could explain why SRRM2 promotes the formation of nuclear tau aggregates.

Our results suggest that the rate limiting step in nuclear tau aggregation is either the production of secondary seeds or their nuclear entry. This is suggested by the observations that not all cells develop nuclear tau aggregates in the time frame of our experiments, but cells with nuclear aggregates typically form multiple separate tau aggregates in a similar time frame (Fig. 2). Strikingly, all nuclear aggregates we observe are localized to nuclear speckles highlighting that these assemblies serve, much as cytoplasmic speckles [2], as preferred sites of tau aggregation. Interestingly, for tau to form nuclear aggregates, it would imply that some monomeric tau can enter the nucleus, which has been suggested previously [41,42].

Our results argue that tau aggregates in the nucleus alter the function of nuclear speckles. This was first suggested by observations that the spatial organization of nuclear speckles was disrupted from a homogeneous overlap of nuclear speckle components into a bipartite assembly with aggregated tau and SRRM2 in the center, and a surrounding region known to contain MSUT2 and poly(A)+ RNA ([5]). Moreover, as assessed by FRAP, nuclear tau aggregates reduce the dynamics of SRRM2 and SRSF2, and possibly other components [5]. Finally, as we show here, cells with nuclear tau aggregates accumulate some specific RNAs and poly(A)+ RNA in the nucleus (Fig. 5). Thus, the aggregation of tau in the nucleus leads to at least alterations in RNA export and possibly could alter pre-mRNA processing [5].

A key question is whether nuclear tau aggregates impact the pathology in human disease. Since nuclear tau aggregates are be observed in cell line and mouse models of disease [5,13], and can be observed in some human patients [15–18], it is possible that such nuclear aggregates could alter nuclear RNA export and/or RNA processing in neurons and thereby contribute to human pathology.

## Acknowledgements

Data analysis of FISH images was performed at the BioFrontiers Institute’s Advanced Light Microscopy Core (RRID: SCR_018302). The Analysis Workstation and the software package Imaris were supported by NIH 1S10RR026680-01A1. Cell culture was performed in the Biochemistry Cell Culture Facility (RRID:SCR_018988). We thank Joseph Dragavon of the BioFrontiers Institute’s Advanced Light Microscopy Core for all of his support for microscopy operating issues and Theresa Nahreini and Emily Proksch for their expert support of cell culture.

## Funding

This work was supported by funds to R.P. from the Howard Hughes Medical Institute.

## Author contributions

C.D., K.M., E.L., J.P., M.V.A., Y.W., and R.P. designed research; C.D., E.L., J.P., M.V.A., and Y.W., performed research and analyzed data. C.D., K.M., J.P., Y.W. and R.P. wrote the paper.

## Ethics declarations

Ethics approval and consent to participate – Not applicable.

Consent for publication – Not applicable.

Competing interests – The authors declare no competing interest.

## Notes

### Competing Interest Statement

The authors have declared no competing interest.

## References

[1] Lee VMY, Goedert M, Trojanowski JQ. Neurodegenerative tauopathies. Annu Rev Neurosci 2001;24. 10.1146/annurev.neuro.24.1.1121.

[2] Lester E, Van Alstyne M, McCann KL, Reddy S, Cheng LY, Kuo J, et al. Cytosolic condensates rich in polyserine define subcellular sites of tau aggregation. Proc Natl Acad Sci U S A 2023;120. 10.1073/pnas.2217759120.

[3] Van Alstyne M, Pratt J, Parker R. Diverse influences on tau aggregation and implications for disease progression. Genes Dev 2025;39:555–81. 10.1101/gad.352551.124.

[4] Ghag G, Bhatt N, Cantu D V., Guerrero-Munoz MJ, Ellsworth A, Sengupta U, et al. Soluble tau aggregates, not large fibrils, are the toxic species that display seeding and cross-seeding behavior. Protein Science 2018;27. 10.1002/pro.3499.

[5] Lester E, Ooi FK, Bakkar N, Ayers J, Woerman AL, Wheeler J, et al. Tau aggregates are RNA-protein assemblies that mislocalize multiple nuclear speckle components. Neuron 2021;109. 10.1016/j.neuron.2021.03.026.

[6] McMillan PJ, Strovas TJ, Baum M, Mitchell BK, Eck RJ, Hendricks N, et al. Pathological tau drives ectopic nuclear speckle scaffold protein SRRM2 accumulation in neuron cytoplasm in Alzheimer’s disease. Acta Neuropathol Commun 2021;9. 10.1186/s40478-021-01219-1.

[7] Jiang L, Lin W, Zhang C, Ash PEA, Verma M, Kwan J, et al. Interaction of tau with HNRNPA2B1 and N6-methyladenosine RNA mediates the progression of tauopathy. Mol Cell 2021;81. 10.1016/j.molcel.2021.07.038.

[8] Shan L, Li P, Yu H, Chen LL. Emerging roles of nuclear bodies in genome spatial organization. Trends Cell Biol 2024;34. 10.1016/j.tcb.2023.10.012.

[9] Faber GP, Nadav-Eliyahu S, Shav-Tal Y. Nuclear speckles – a driving force in gene expression. J Cell Sci 2022;135. 10.1242/jcs.259594.

[10] Ferreira JA, Carmo-Fonseca M, Lamond AI. Differential interaction of splicing snRNPs with coiled bodies and interchromatin granules during mitosis and assembly of daughter cell nuclei. Journal of Cell Biology 1994;126. 10.1083/jcb.126.1.11.

[11] Van Alstyne M, Brown G, Nguyen VL, Ramaswami M, Hoeffer CA, Parker R. Polyserine-mediated targeting of FAF2/UBXD8 ameliorates tau aggregation. Neuron 2025;113. 10.1016/j.neuron.2025.08.002.

[12] Pratt J, McCann K, Kuo J, Parker R. Polyserine-tau interactions modulate tau fibrillization. J Biol Chem 2025;301:110523. 10.1016/j.jbc.2025.110523.

[13] Sanders DW, Kaufman SK, DeVos SL, Sharma AM, Mirbaha H, Li A, et al. Distinct tau prion strains propagate in cells and mice and define different tauopathies. Neuron 2014;82. 10.1016/j.neuron.2014.04.047.

[14] Kang SG, Han ZZ, Daude N, McNamara E, Wohlgemuth S, Molina-Porcel L, et al. Pathologic tau conformer ensembles induce dynamic, liquid-liquid phase separation events at the nuclear envelope. BMC Biol 2021;19. 10.1186/s12915-021-01132-y.

[15] Bengoa-Vergniory N, Velentza-Almpani E, Silva AM, Scott C, Vargas-Caballero M, Sastre M, et al. Tau-proximity ligation assay reveals extensive previously undetected pathology prior to neurofibrillary tangles in preclinical Alzheimer’s disease. Acta Neuropathol Commun 2021;9. 10.1186/s40478-020-01117-y.

[16] Metuzals J, Robitaille Y, Houghton S, Gauthier S, Leblanc R. Paired helical filaments and the cytoplasmic-nuclear interface in Alzheimer’s disease. J Neurocytol 1988;17. 10.1007/BF01216709.

[17] Montalbano M, McAllen S, Puangmalai N, Sengupta U, Bhatt N, Johnson OD, et al. RNA-binding proteins Musashi and tau soluble aggregates initiate nuclear dysfunction. Nat Commun 2020;11. 10.1038/s41467-020-18022-6.

[18] Papasozomenos Ch. S. Nuclear tau immunoreactivity in presenile dementia with motor neuron disease: A case report. Clin Neuropathol 1995;14.

[19] Hochmair J, Exner C, Franck M, Dominguez-Baquero A, Diez L, Brognaro H, et al. Molecular crowding and RNA synergize to promote phase separation, microtubule interaction, and seeding of Tau condensates. EMBO J 2022;41. 10.15252/embj.2021108882.

[20] Saha I, Yuste-Checa P, Da Silva Padilha M, Guo Q, Körner R, Holthusen H, et al. The AAA+ chaperone VCP disaggregates Tau fibrils and generates aggregate seeds in a cellular system. Nat Commun 2023;14. 10.1038/s41467-023-36058-2.

[21] Batra S, Vaquer-Alicea J, Valdez C, Taylor SP, Manon VA, Vega AR, et al. VCP regulates early tau seed amplification via specific cofactors. Molecular Neurodegeneration 2025;20. 10.1186/s13024-024-00783-z.

[22] Ariotti N, Rae J, Giles N, Martel N, Sierecki E, Gambin Y, et al. Ultrastructural localisation of protein interactions using conditionally stable nanobodies. PLoS Biol 2018;16. 10.1371/journal.pbio.2005473.

[23] Khong A, Matheny T, Jain S, Mitchell SF, Wheeler JR, Parker R. The Stress Granule Transcriptome Reveals Principles of mRNA Accumulation in Stress Granules. Mol Cell 2017;68. 10.1016/j.molcel.2017.10.015.

[24] Gaspar I, Wippich F, Ephrussi A. Enzymatic production of single-molecule FISH and RNA capture probes. RNA 2017;23. 10.1261/rna.061184.117.

[25] Ilık İA, Malszycki M, Lübke AK, Schade C, Meierhofer D, Aktaş T. Son and srrm2 are essential for nuclear speckle formation. Elife 2020;9. 10.7554/eLife.60579.

[26] Holmes BB, Furman JL, Mahan TE, Yamasaki TR, Mirbaha H, Eades WC, et al. Proteopathic tau seeding predicts tauopathy in vivo. Proc Natl Acad Sci U S A 2014;111. 10.1073/pnas.1411649111.

[27] Kang SG, Eskandari-Sedighi G, Hromadkova L, Safar JG, Westaway D. Cellular Biology of Tau Diversity and Pathogenic Conformers. Front Neurol 2020;11. 10.3389/fneur.2020.590199.

[28] Braak H, Braak E. CORTICAL AND SUBCORTICAL ARGYROPHILIC GRAINS CHARACTERIZE A DISEASE ASSOCIATED WITH ADULT ONSET DEMENTIA. Neuropathol Appl Neurobiol 1989;15. 10.1111/j.1365-2990.1989.tb01146.x.

[29] Tolnay M, Spillantini MG, Goedert M, Ulrich J, Langui D, Probst A. Argyrophilic grain disease: Widespread hyperphosphorylation of tau protein in limbic neurons. Acta Neuropathol 1997;93. 10.1007/s004010050642.

[30] Eftekharzadeh B, Daigle JG, Kapinos LE, Coyne A, Schiantarelli J, Carlomagno Y, et al. Tau Protein Disrupts Nucleocytoplasmic Transport in Alzheimer’s Disease. Neuron 2018;99. 10.1016/j.neuron.2018.07.039.

[31] Woerner AC, Frottin F, Hornburg D, Feng LR, Meissner F, Patra M, et al. Cytoplasmic protein aggregates interfere with nucleocytoplasmic transport of protein and RNA. Science (1979) 2016;351. 10.1126/science.aad2033.

[32] Dasso M, Pu RT. Nuclear transport: Run by ran? Am J Hum Genet 1998;63. 10.1086/301990.

[33] Dworak N, Makosa D, Chatterjee M, Jividen K, Yang CS, Snow C, et al. A nuclear lamina-chromatin-Ran GTPase axis modulates nuclear import and DNA damage signaling. Aging Cell 2019;18. 10.1111/acel.12851.

[34] Grima JC, Daigle JG, Arbez N, Cunningham KC, Zhang K, Ochaba J, et al. Mutant Huntingtin Disrupts the Nuclear Pore Complex. Neuron 2017;94. 10.1016/j.neuron.2017.03.023.

[35] Zhang K, Donnelly CJ, Haeusler AR, Grima JC, Machamer JB, Steinwald P, et al. The C9orf72 repeat expansion disrupts nucleocytoplasmic transport. Nature 2015;525. 10.1038/nature14973.

[36] Anderson DJ, Le Moigne R, Djakovic S, Kumar B, Rice J, Wong S, et al. Targeting the AAA ATPase p97 as an Approach to Treat Cancer through Disruption of Protein Homeostasis. Cancer Cell 2015;28. 10.1016/j.ccell.2015.10.002.

[37] Magnaghi P, D’Alessio R, Valsasina B, Avanzi N, Rizzi S, Asa D, et al. Covalent and allosteric inhibitors of the ATPase VCP/p97 induce cancer cell death. Nat Chem Biol 2013;9. 10.1038/nchembio.1313.

[38] Masuda S, Das R, Cheng H, Hurt E, Dorman N, Reed R. Recruitment of the human TREX complex to mRNA during splicing. Genes Dev 2005;19. 10.1101/gad.1302205.

[39] Wang K, Wang L, Wang J, Chen S, Shi M, Cheng H. Intronless mRNAs transit through nuclear speckles to gain export competence. Journal of Cell Biology 2018;217. 10.1083/jcb.201801184.

[40] Candia RF, Cohen LS, Morozova V, Corbo C, Alonso AD. Importin-Mediated Pathological Tau Nuclear Translocation Causes Disruption of the Nuclear Lamina, TDP-43 Mislocalization and Cell Death. Front Mol Neurosci 2022;15. 10.3389/fnmol.2022.888420.

[41] Younas N, Saleem T, Younas A, Zerr I. Nuclear face of Tau: an inside player in neurodegeneration. Acta Neuropathol Commun 2023;11. 10.1186/s40478-023-01702-x.

[42] Antón-Fernández A, Vallés-Saiz L, Avila J, Hernández F. Neuronal nuclear tau and neurodegeneration. Neuroscience 2023;518. 10.1016/j.neuroscience.2022.07.015.

