## Supplemental materials for "Nuclear tau aggregates inhibit RNA export and form by secondary seeding from cytosolic tau aggregates"

**List of Supplementary Materials**

Figs. S1 to S6

Movies S1 to S4

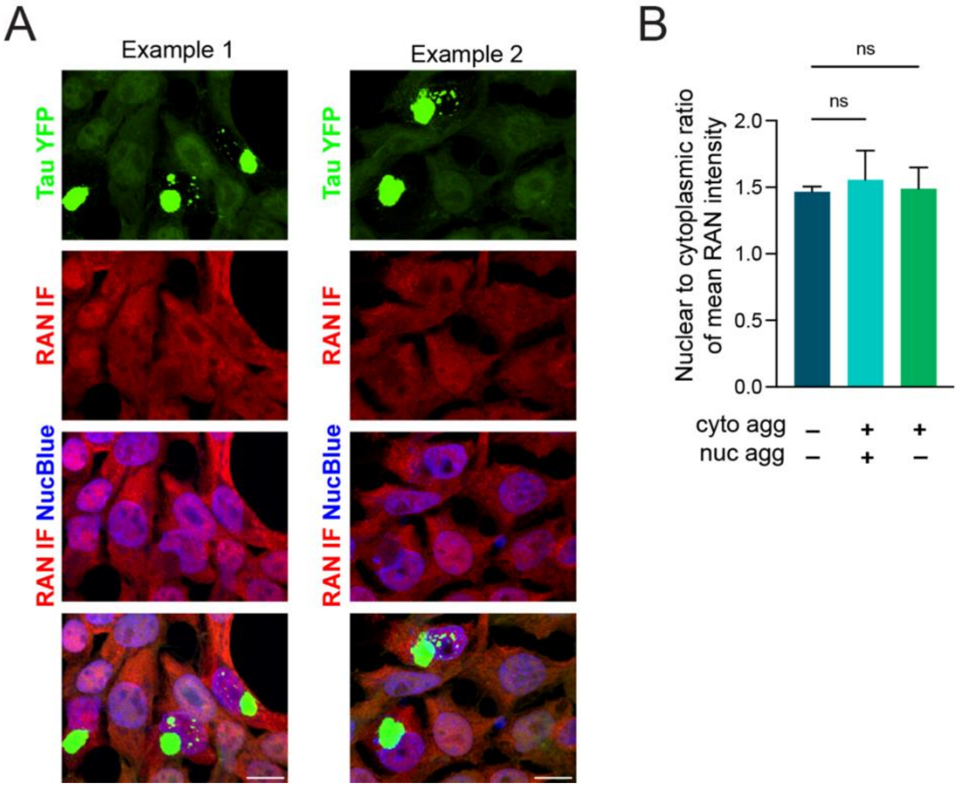


**Fig. S1 Nuclear to cytoplasmic ratio of RAN in cells with or without tau aggregates in the nucleus or cytoplasm.** (**A**) Two examples of RAN immunofluorescence in HEK293T tau biosensor cells with no tau aggregates, cytoplasmic aggregates only, or both nuclear and cytoplasmic aggregates. (**B**) Nuclear to cytoplasmic ratio of RAN in cells with or without tau aggregates. Bar for all plots represent mean and SD. N = 3 independent experiments, statistics performed with ordinary one-way Anova and Dunnett's multiple comparisons, ns = not significant.

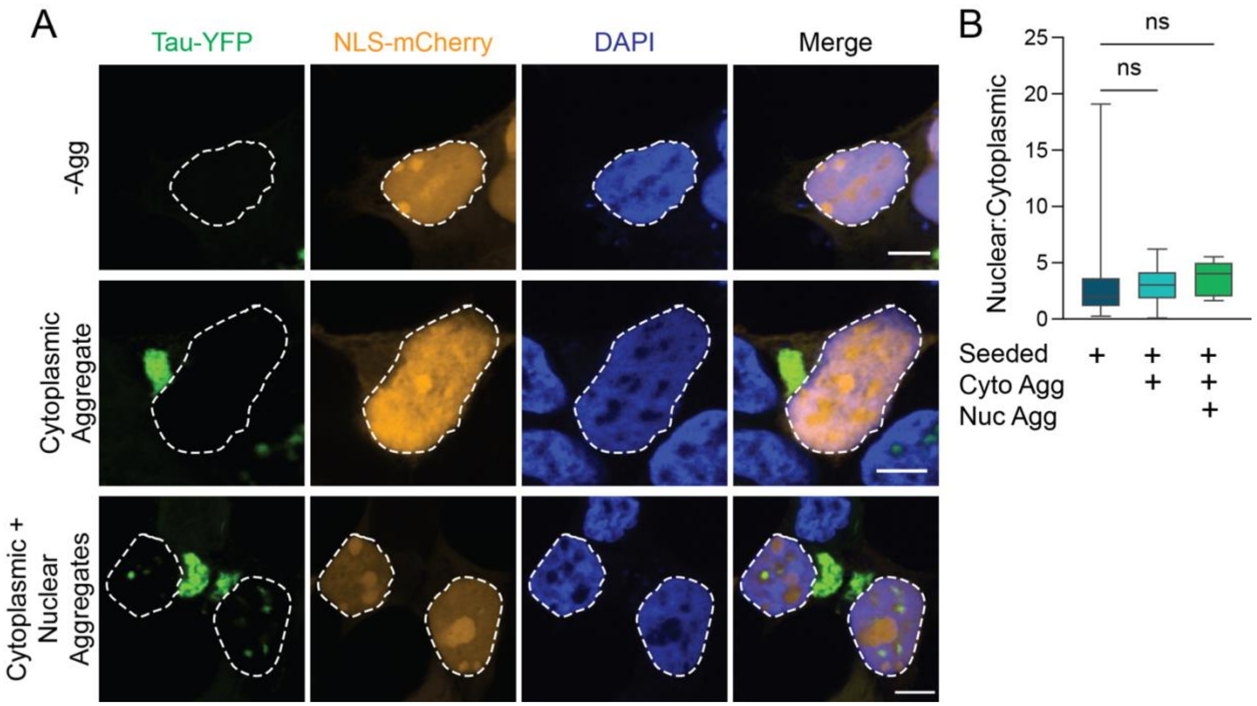


**Fig. S2 Tau aggregation does not alter nuclear envelope integrity or nuclear import activity.** (**A**) Representative images of cells transfected with a plasmid encoding NLS-mCherry with and without tau aggregation in the nucleus or cytoplasm. White outlines show nuclear envelope periphery from DAPI channel, overlaid to other channels. (**B**) Quantification of the average nuclear:cytoplasmic ratio of NLS-mCherry from single cells. N = 3 biological replicates, statistics performed with ordinary one-way ANOVA and Dunnett's multiple comparisons test, ns = non-significant.

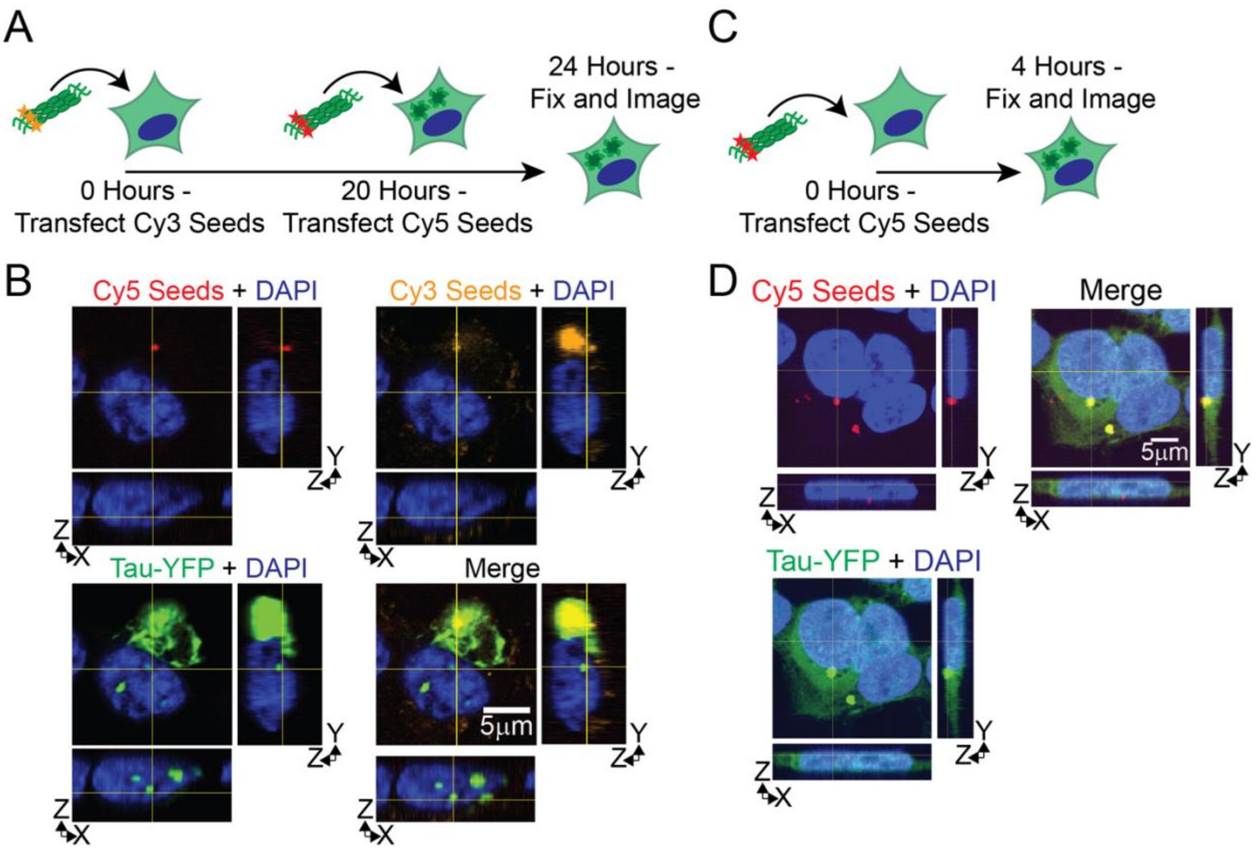


**Fig. S3 Exogenous tau seeds are not in nuclear aggregates with multiple seeding events** (**A**) Experiment schematic where Cy3 tau seeds are transfected into HEK293T tau biosensor cells, followed by transfection of Cy5 labeled tau seeds 20 hours later. Cells were fixed 24 hours after transfection of Cy3 seeds. (**B**) Representative images of seeding experiment described in (A). Square images are a single Z plane in XY, YZ and XZ projections are shown to the right and below for each channel. (**C**) Experiment schematic where Cy5 labeled tau seeds are transfected into HEK293T tau biosensor cells and fixed 4 hours later. (**D**) Representative images of experiment described in (C). Square images are a single Z plane in XY, YZ and XZ projections are shown below and to the right of each channel.

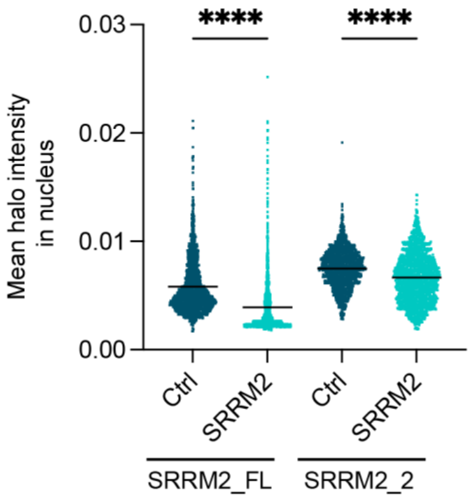


**Fig. S4 DsiRNA knockdown of polyserine domain containing SRRM2 reduces the intensity of Halo in the nucleus**. Graph of the mean halo intensity in the nuclei of SRRM2_FL halo and SRRM2_2 halo HEK293T tau biosensor cells after treatment with either a control DsiRNA or a DsiRNA against polyserine domain absent from SRRM2_2. Bar indicates mean. N = 3 independent experiments, statistics performed using ordinary one-way Anova and Sidak's multiple comparisons test, **** = $p < 0.0001$.

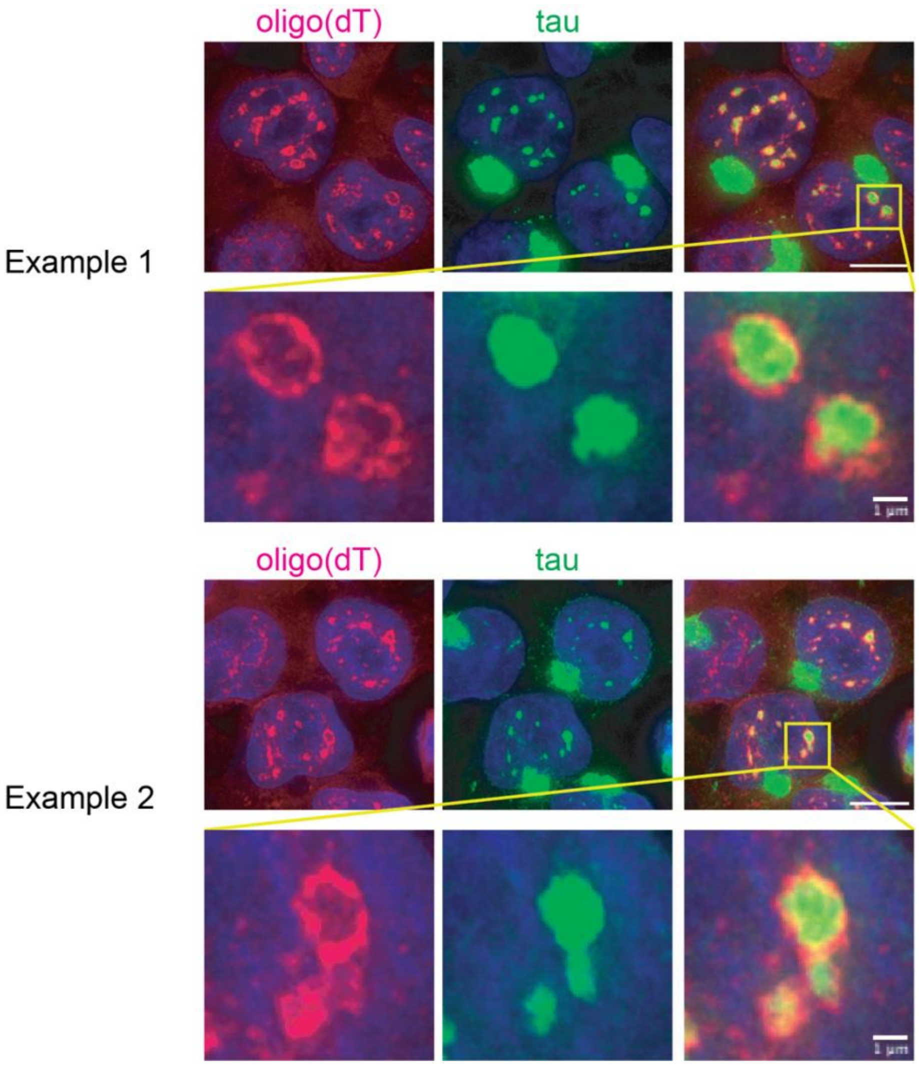


**Fig. S5 PolyA+ FISH signal forms a periphery around nuclear tau aggregates.** Representative images of FISH with oligo(dT) probes in HEK293T tau biosensor cells treated with tau mouse brain homogenate.

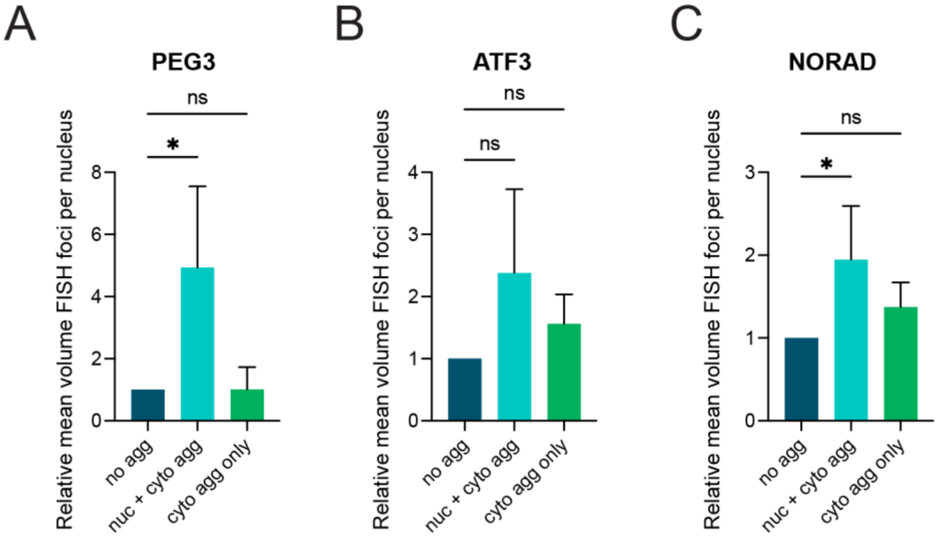


**Fig. S6 The presence of nuclear tau aggregates alters the volume of FISH foci in the nucleus.** (**A-C**) The mean volume of FISH foci for the indicated RNAs in cells with only cytoplasmic aggregates or with both nuclear and cytoplasmic aggregates relative to the mean number observed in cells with no aggregates. Bars for all plots represent mean and SD. (**A**) Mean volume of PEG3 FISH foci. N = 3 independent experiments. Statistics performed with ordinary one-way Anova and Dunnett's multiple comparisons test, ns = not significant, * p = 0.0379. (**B**) Mean volume of ATF3 FISH foci. N =. 3 independent experiments. Statistics performed with ordinary one-way Anova and Dunnett's multiple comparisons test, ns = not significant. (**C**) Mean volume of NORAD FISH foci. N = 4 independent experiments. Statistics performed with ordinary one-way Anova and Dunnett's multiple comparisons test, ns = not significant, * p = 0.0184.

**Movie S1.** Movie showing nuclear tau aggregation (tau-YFP, green) following mitosis. Imaging initiated 6 hours after tau seeding.

**Movie S2**. Movie showing nuclear tau aggregates (tau-YFP, green) forming in cells following cytoplasmic tau aggregation and nuclear ring formation without going through mitosis. Imaging initiated 6 hours after tau seeding. Example 1.

**Movie S3.** Movie showing nuclear tau aggregates (tau-YFP, green) forming in cells following cytoplasmic tau aggregation and nuclear ring formation without going through mitosis. Imaging initiated 6 hours after tau seeding. Example 2.

**Movie S4.** Movie showing nuclear tau aggregates (tau-YFP, green) forming in cells following cytoplasmic tau aggregation and nuclear ring formation without going through mitosis. Imaging initiated 0.5 hours after tau seeding. Example 3.
